# TumorArchetypeR: A modular framework to derive signature-based tumor subtypes

**DOI:** 10.64898/2026.05.11.724259

**Authors:** Mechthild Lütge, Sina Nassiri

**Affiliations:** Roche Innovation Center Zurich, Wagistrasse 10, Schlieren 8952, Switzerland; Roche Innovation Center Basel, Grenzacherstrasse 124, 4070 Basel, Switzerland

## Abstract

**Motivation:** The tumor microenvironment (TME) dictates cancer progression and therapeutic response, yet translating TME subtypes into robust clinical biomarkers remains a significant challenge. Existing classification models typically rely on static gene signatures and cohort-dependent normalization, making them ill-suited for application to the small, unbalanced datasets common in early-phase clinical trials. To better guide drug development, methods are required that offer the flexibility to target specific biological contexts and bridge the gap between the discovery of tumor archetypes and their robust translation to individual patient samples.

**Results:** We developed TumorArchetypeR, a modular R package that unifies unsupervised subtype discovery with the generation of rank-based, single-sample classifiers. By leveraging a systematic parameter grid search, the framework identifies stable, data-driven subtypes rather than relying on arbitrary defaults. Crucially, to ensure clinical translatability, the package includes a module to train a robust classifier using binary gene-pair rules, enabling prediction without cohort-level preprocessing. Applying TumorArchetypeR to colorectal cancer, we resolved the heterogeneity of fibrotic tumors, distinguishing an immunosuppressive “Immune-enriched/Fibrotic” state from an immune-excluded “Fibrotic/Myeloid” phenotype. Furthermore, we identified a distinct “Th/B-cell enriched” archetype associated with superior survival, a group largely obscured by existing pan-cancer models. With our rank-based classifier demonstrating robust performance on previously unseen samples, these findings highlight TumorArchetypeR as a scalable, end-to-end solution for refining patient stratification and optimizing precision oncology strategies. The TumorArchetypeR package and documentation are openly available on GitHub at https://github.com/lutgem/TumorArchetypeR.

## Introduction

The tumor microenvironment (TME) fundamentally shapes cancer progression, therapeutic efficacy, and patient survival. Consequently, identifying ‘tumor archetypes’, defined as recurrent, biologically distinct cellular compositions or states within the tumor ecosystem, has emerged as a powerful, biologically driven framework for classifying solid tumors, offering the potential to guide drug development from early-phase discovery to clinical application (Thorsson et al. 2018; Combes, Samad and Krummel 2023). Emerging pan-cancer studies have established that TME-based subtyping provides patient stratification beyond classical genomic or histological classifications (Combes et al. 2022; Bagaev et al. 2021). For instance, Bagaev et al. identified four conserved TME states, ranging from immune-depleted to immune-enriched and fibrotic, that predict immunotherapy response and survival across diverse tumor types, underscoring the immense potential of resolving microenvironmental heterogeneity to inform drug development processes (Bagaev et al. 2021).

Despite this potential, existing archetype models suffer from fundamental limitations that hinder their adaptability and clinical translation. First, established classification systems are typically “static,” relying on fixed gene signatures defined at the time of their publication. These rigid models cannot easily incorporate evolving biological knowledge, such as newly discovered cell states derived from single-cell RNA sequencing or novel therapeutic targets. Second, the computational approaches used to derive these archetypes are typically cohort-dependent. They require large, representative datasets for preprocessing and normalization, making them ill-suited for application to independent datasets, small cohorts, or individual patient samples common in early-phase drug development.

To address these challenges, we developed TumorArchetypeR, a flexible computational framework designed to bridge the gap between the *de novo* discovery of biologically driven tumor subtypes and their robust translation to unseen data. Unlike static models, TumorArchetypeR provides a modular pipeline that enables researchers to define archetypes based on any user-provided set of biological signatures, facilitating the integration of diverse molecular features. The framework leverages a systematic parameter grid search to explore the stability and biological relevance of multiple competing models, moving beyond arbitrary default parameters toward data-driven model selection. Crucially, to address the challenge of clinical translatability, the framework includes a dedicated module for training robust, rank-based classifiers. By utilizing binary gene-pair rules that rely solely on within-sample relative ordering, these classifiers generate predictions that are stable across platforms and applicable to individual samples without the need for cohort-based preprocessing.

In this study, we demonstrate the utility of TumorArchetypeR by applying it to transcriptomic data from the TCGA colorectal cancer cohort. We derived a refined TME subtyping model that resolves the heterogeneity of fibrotic tumors, distinguishing between immunosuppression and immune exclusion. We further characterize the genomic underpinnings of these archetypes, linking distinct immune landscapes to specific mutational drivers such as *APC*-driven WNT signaling and the serrated neoplasia pathway. Our findings highlight TumorArchetypeR as a powerful, end-to-end solution for identifying reproducible tumor archetypes and translating them to refine patient stratification and optimizing precision oncology strategies.

## Results

### TumorArchetypeR: A flexible framework to derive signature-based tumor subtypes

Existing signature-based tumor subtyping models suffer from a lack of flexibility and limited translatability to previously unseen data. To address these limitations we developed TumorArchetypeR, a modular R package designed to bridge the gap between targeted discovery of *de novo* subtypes and robust translation across datasets. The framework is built upon three major steps: (1) the systematic derivation of archetype models via parameter grid search, (2) data-driven model selection, and (3) classifier training for translation to new datasets (Fig. 1).

**Figure 1.**
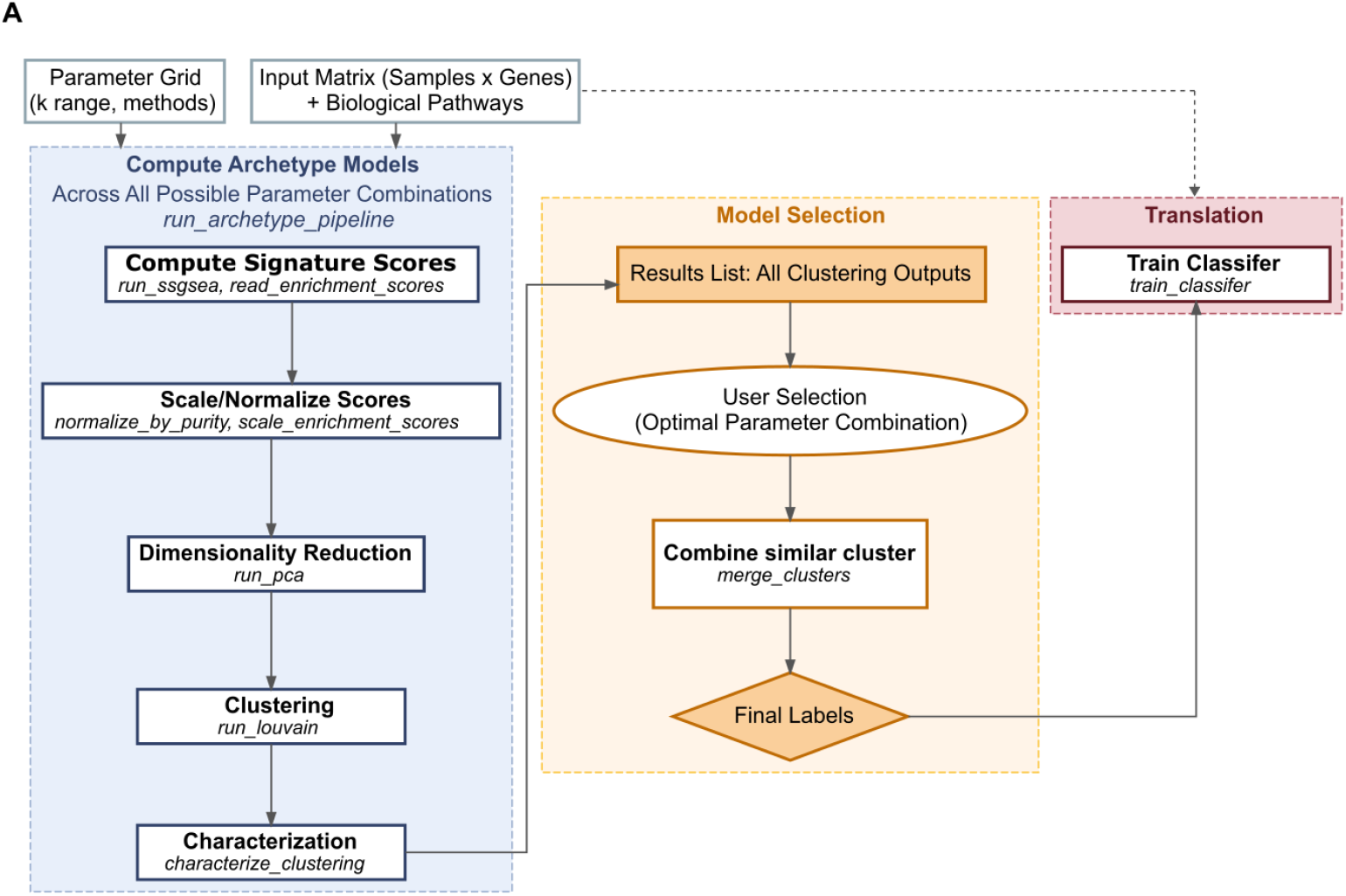
Schematic overview of the TumorArchetypeR framework. The workflow is structured into three modular phases designed to bridge *de novo* subtype discovery with robust clinical translation. (Phase 1) Systematic Archetype Discovery. The framework accepts either raw gene expression data or pre-computed enrichment scores as input. The wrapper function *run_archetype_pipeline* orchestrates a parallel grid search across a user-defined parameter space and executes a sequential pipeline comprising signature scoring, data scaling/normalization, dimensionality reduction, clustering, and characterization. (Phase 2) Model Selection. The User can select the optimal subtyping model guided by a number of clustering stability metrics, visualization functions, and associations with clinical covariates. Optionally the model can be further refined by merging similar clusters. (Phase 3) Classifier Training and Translation. The final module utilized rank-based binary gene-pair rules derived from the raw gene expression data to train a random forest classifier on the selected model. By using within-sample relative ordering the classifier ensures robust subtype assignment in independent datasets, distinguishing it from cohort-dependent methods and enabling application to single samples.

Input data for the framework can be either a raw gene expression matrix coupled with biological signatures of interest or pre-computed enrichment scores. From this input, archetype models are generated through a sequential pipeline comprising: (1) signature scoring, (2) scaling and/or normalization, (3) dimensionality reduction, (4) clustering, and (5) cluster characterization. This entire sequence is orchestrated by a single wrapper function, “run_archetype_pipeline”, which runs across all possible combinations of a user defined parameter grid. Efficient parallel execution is facilitated by utilizing the pipeComp framework, which ensures the caching of intermediate results and avoids redundant computations (Germain, Sonrel and Robinson 2020). A systematic exploration of models derived from the different parameter configurations is supported through the calculation of stability metrics, covariate association and multiple visualization functions, ensuring the identification of an optimal, robust model rather than relying on arbitrary default parameters. Finally, to facilitate a seamless translation of the discovered subtypes to independent datasets, the framework provides a module to train a robust classifier from either gene expression data or signature scores. Importantly, because the classifier utilizes rank-based binary gene-pair rules, it relies on within-sample relative ordering, rendering predictions independent of cohort-level normalization and applicable to single samples or unbalanced cohorts (Marzouka and Eriksson 2021).

### Application of the TumorArchetypeR framework identifies TME subtypes in colorectal cancer

To demonstrate the practical utility and robustness of TumorArchetypeR, we applied the framework to derive Tumor Microenvironment (TME) subtypes using transcriptomic data from the TCGA Colorectal Cancer cohort (N = 378 patients). To avoid data leakage during classifier validation, the cohort was randomly partitioned into a discovery set (80%), used to derive the subtyping model and train the classifier, and a held-out validation set (20%), reserved to assess the translatability of the final model.

For initial feature generation, we utilized xCell2.0, a state-of-the-art, enrichment-based cell type deconvolution tool (Angel et al. 2025). To capture the full heterogeneity of the TME, we calculated enrichment scores using two complementary references available in the xCell2.0 repository: (1) the Tumor Microenvironment reference (TMEref) and (2) the Pan-Cancer Immune reference (PanCref). Initial feature exploration revealed a high degree of collinearity across the enrichment scores, identifying distinct clusters corresponding to major TME cell lineages (Supp. Fig. 1A). In addition we observed a substantial heterogeneity in score distributions, with marked differences in both mean signal intensity and variance across cell type scores (Supp. Fig. 1B). To identify robust subtypes from this heterogenous data, we executed the TumorArchetypeR pipeline on the discovery set. The workflow comprised feature scaling as an initial step to prevent highly abundant cell populations (e.g., malignant cells) from dominating the clustering structure. Crucially, to prevent the artificial amplification of uninformative noise during scaling, the pipeline incorporates a variance-based filtering step based on a user-defined quantile threshold. Accordingly, we excluded the 13 features with the lowest variance, thereby removing low-signal signatures such as NK cell, mast cell, and glial cell scores. To further mitigate noise and resolve feature redundancy, the pipeline subsequently applied dimensionality reduction via Principal Component Analysis (PCA) prior to Louvain clustering on a nearest-neighbor graph. Analysis of feature loadings confirmed that the top dimensions effectively condensed the input space, with each PC capturing a distinct major axis of TME variation (Supp. Fig. 1C).

A defining feature of the TumorArchetypeR framework is its capacity to simultaneously derive multiple competing models through an efficient, parallel parameter grid search. For this application, we defined the grid to systematically explore two critical hyperparameters: a range of cumulative variance thresholds (determining the number of input PCs) (Supp Fig. 1D) and varying k-nearest neighbors for Louvain graph generation. To guide the identification of the optimal model, TumorArchetypeR employs a multi-faceted evaluation strategy. First, it computes a suite of structure-based validity indices (including Silhouette width, Calinski-Harabasz Index, and Dunn Index) alongside a clustering stability estimate derived from the mean Adjusted Rand Index (ARI) of repeated bootstrap runs. Recognizing that individual metrics have specific limitations, the pipeline synthesizes these inputs into a single Combined Clustering Score. This composite metric is calculated as the arithmetic mean of the normalized structure-based indices and the normalized stability estimate, balancing cluster coherence with robustness (Fig. 2A).

**Figure 2.**
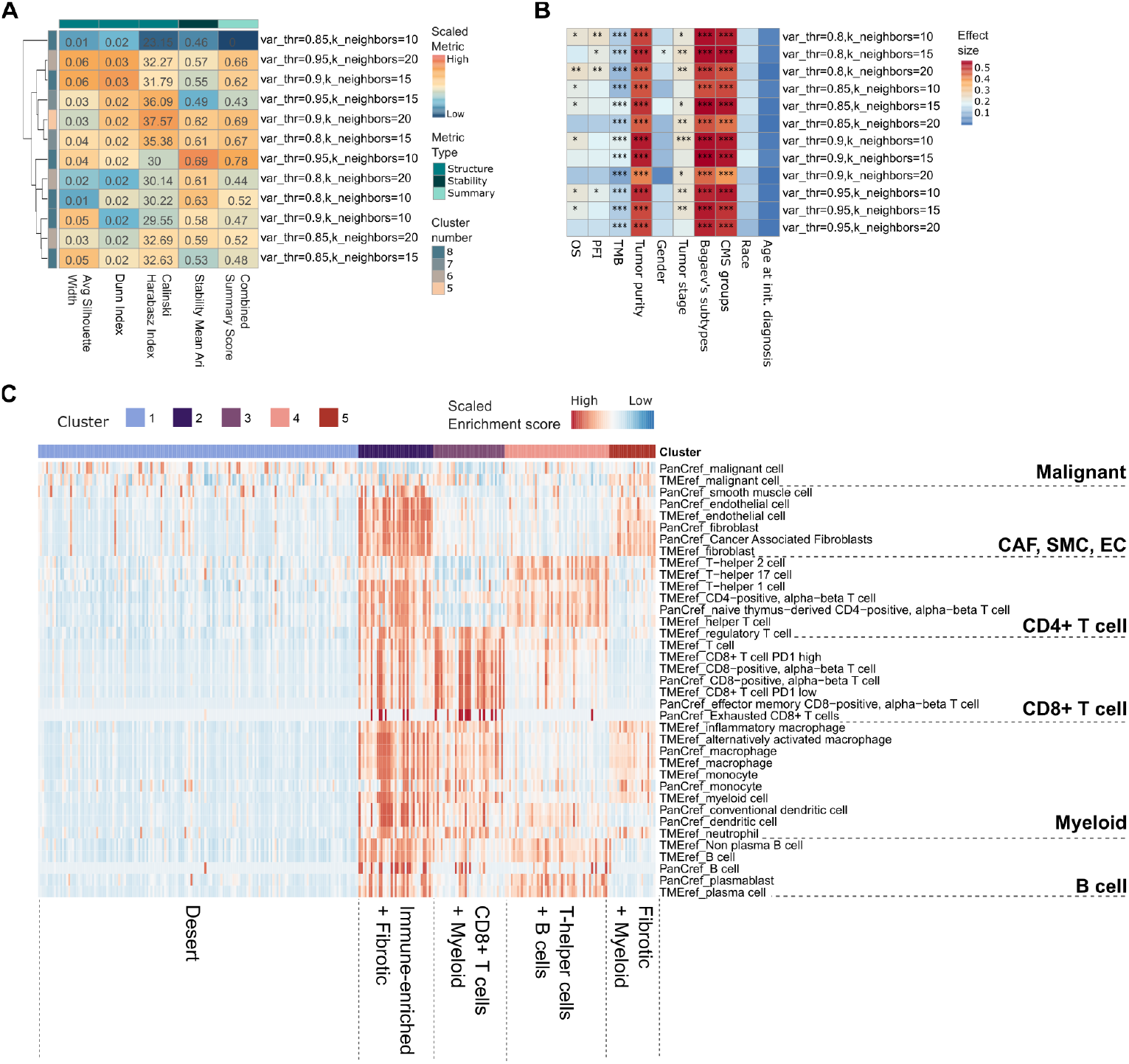
Data-driven model selection of TME archetypes. (**A**) Systematic evaluation of clustering models across the parameter grid. Heatmap showing the scaled clustering metrics alongside the composite Combined Summary Score across all executed parameter combinations. Rows represent individual parameter runs, ordered by hierarchical clustering. Numbers display the unscaled values for each clustering metric and the Combined Summary Score. (**B**) Assessment of clinical and biological relevance. Heatmap displaying the association of each clustering model (rows) with user-provided covariates (columns). Heatmap colors represents the effect size, calculated specifically for each data type: Cramer’s V for categorical variables, Epsilon-squared for numeric variables, and a Phi-like metric 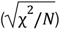 for survival endpoints. Asterisks indicate the statistical significance level derived from log-rank tests (survival) or covariate association tests (Chi-squared/Kruskal-Wallis). (C) Heatmap displaying the distinct enrichment score profiles of the five archetypes identified following automated selection and cluster merging.

Second, beyond internal validation, the pipeline evaluates biological and clinical relevance by calculating effect sizes and significance estimates for user-provided covariates and survival endpoints (Fig. 2B). To facilitate this, the user provides sample metadata and specifies relevant clinical variables. For each survival endpoint, the pipeline generates Kaplan-Meier curves and performs log-rank tests. For numeric covariates, associations are assessed via Kruskal-Wallis tests, while categorical variables are evaluated using chi-squared tests. These quantitative outputs are complemented by a suite of visualization functions for inspecting metric distributions, covariate associations, and enrichment scores. In the discovery dataset, analysis revealed strong, highly significant associations with tumor mutational burden (TMB), tumor purity, Bagaev’s TME subtypes (Bagaev et al. 2021), and Consensus Molecular Subtypes (CMS) groups (Guinney et al. 2015) across all clustering runs, while no significant associations were found for age or race. Interestingly, only a subset of clustering runs showed a significant association with overall survival (OS) or progression-free interval (PFI). While the framework supports manual inspection, it defaults to automated selection based on the maximal Combined Clustering Score. Notably, the optimal clustering run (variance threshold = 0.95, k-neighbors = 10) demonstrated significant associations with both OS and PFI.

Following selection, the pipeline enables optional merging of phenotypically similar clusters through hierarchical clustering of their summary profiles. Users can merge based on centroid profiles (mean enrichment), eigengene profiles (the first principal component, inspired by WGCNA (Langfelder and Horvath 2008)), or k-NN graph connectivity. Similarity is derived via Euclidean distance or Pearson correlation, with the final cluster count determined by a user-defined threshold. For the CRC cohort, we applied the automated selection protocol followed by cluster merging using Euclidean distance on centroid profiles (Supp Fig. 2A+B). The resulting final model yielded five distinct TME archetypes, characterized by their enrichment profiles as: (1) Desert, (2) Immune-enriched/Fibrotic, (3) CD8+ T-cell/Myeloid, (4) Th/B-cell enriched, and (5) Fibrotic/Myeloid (Fig. 2C).

Training a rank-based classifier enables robust translation of TME archetypes to independent samples To ensure the broad applicability and clinical translatability of the derived archetype model, the TumorArchetypeR pipeline includes a dedicated classifier training module. A key strength of this module is its reliance on within-sample relative ordering (rank-based binary gene-pair rules) (Marzouka and Eriksson 2021). By focusing on the relative expression of features rather than absolute values, the classifier achieves high robustness across different platforms and normalization methods, enabling reliable predictions for individual samples or small cohorts—common scenarios in clinical practice and early-phase trials. The classifier allows training on either gene expression data or enrichment scores, offering flexibility when translating to new datasets (no additional preprocessing like normalization, scaling or derivation of enrichment scores required).

To maximize model performance, the module incorporates an optional quality control and optimization step prior to training. This includes a consensus sample filtering process (based on cluster Silhouette scores, k-Nearest Neighbor (kNN) consistencies, and Random Forest Out-of-Bag (OOB) probabilities) to retain only high-confidence samples for rule derivation, alongside an optional class balancing step to prevent bias against rare subtypes (Fig. 3A). At its core, the classifier leverages the Random Forest implementation of the multiClassPairs framework to generate decision rules (Marzouka and Eriksson 2021). Model performance is assessed via k-fold cross-validation on the filtered dataset before the final classifier is trained on the full data.

**Figure 3.**
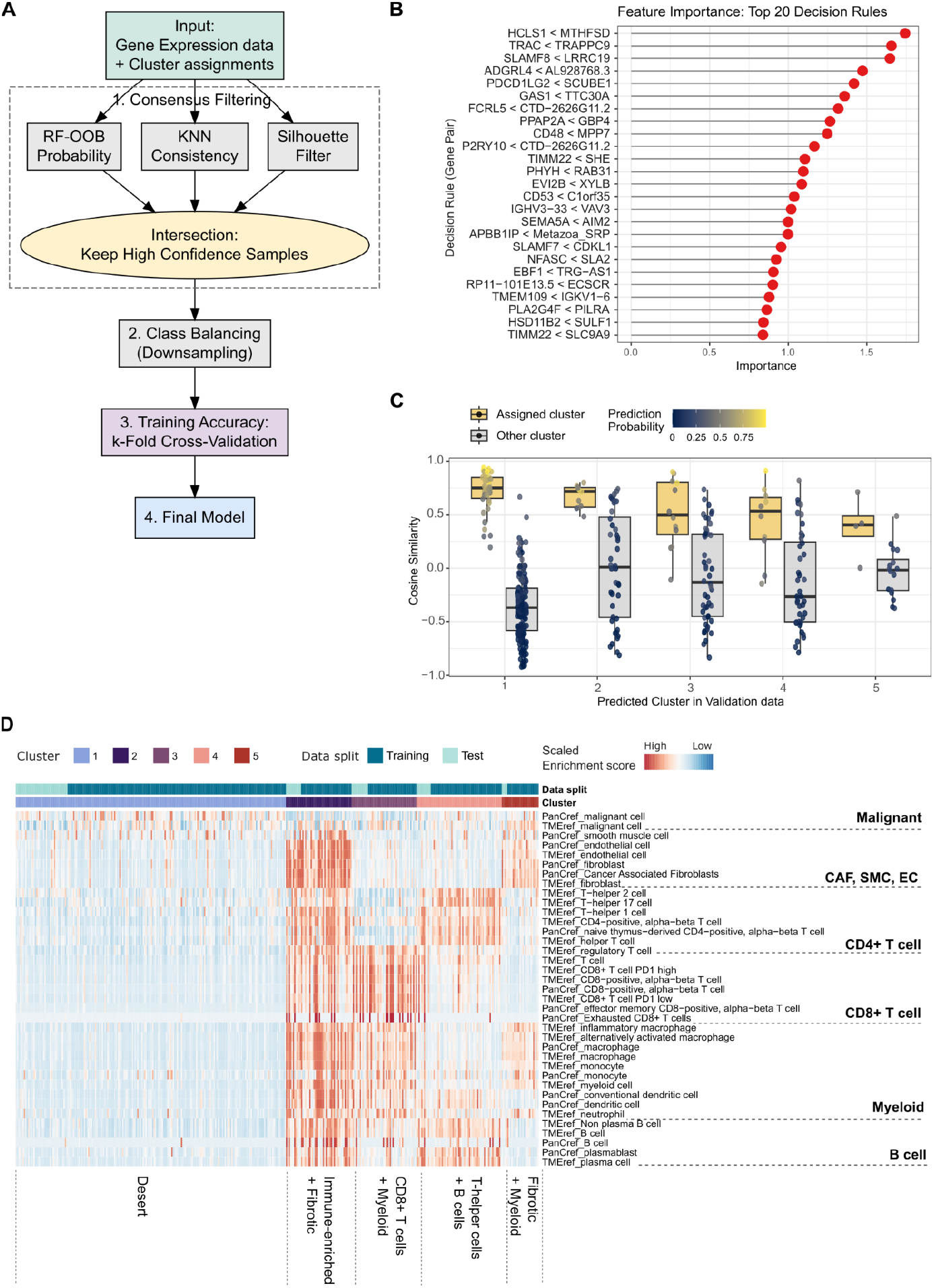
Training and independent validation of a translatable TME subtype classifier. (**A**) Schematic overview of the classifier training module. To maximize robustness, the pipeline incorporates an optional quality control step where samples are consensus-filtered based on Silhouette scores, kNN consistency, and Random Forest Out-of-Bag (OOB) probabilities. An optional class balancing step prevents bias against rare subtypes before final training. (**B**) Characterization of the top discriminative features. The plot displays the top 20 binary gene-pair rules ranked by feature importance. (**C**) Boxplots showing the cosine similarity of samples in the independent validation set (20% of the original cohort) to the centroids of the archetypes derived from the discovery set (80% of the original cohort). Validation samples exhibit higher similarity to their assigned archetype centroid than to centroids from alternative subtypes. Dots represent individual samples and are colored by their classifier prediction probabilities. (**D**) Reproducibility of TME phenotypes. Comparison of enrichment score profiles between the Discovery (training) and Validation (test) cohorts. The validation set fully recapitulates the distinct transcriptional landscapes defined in the discovery set, demonstrating the stability and translatability of the derived model.

When applied to the CRC discovery dataset, the trained classifier demonstrated high performance, achieving an overall training accuracy of 94.5%. Misclassifications were predominantly restricted to the “Desert” cluster (Cluster 1) (Supp Fig. 3A). Notably, these falsely assigned samples consistently exhibited lower classifier-derived class probabilities compared to correctly assigned samples (Supp Fig. 3B), suggesting that they likely represent biological transition states with intermediate immune or fibroblast abundance.

Analysis of the binary gene-pair rule profiles revealed five sample clusters with distinct activation patterns, confirming that the classifier selected highly discriminative rules for each subtype (Supp Fig. 3C). Furthermore, biological interrogation of the top 20 rules (ranked by feature importance) validated that the model successfully captures known TME biology (Fig. 3B). These rules consistently paired known lineage markers, spanning general immune (*CD48, HCLS1*), T-cell (*TRAC, TRG-AS1*), B-cell (*IGHV3-33, FCRL5*), myeloid (*SLAMF8, EVI2B*), and stromal/endothelial (*SULF1, ECSCR*) populations, against ubiquitously expressed housekeeping genes (e.g., *TIMM22, RAB31*) or tumor-intrinsic markers (e.g., *CDKL1*). This specific pairing structure demonstrates that the classifier effectively exploits the relative abundance of TME components versus tumor/background signals to discriminate between subtypes.

To further validate the classifier, we applied the trained model to the independent validation set (20% of the initial cohort), which had been excluded from model derivation and classifier training. We evaluated the phenotypic alignment of these predictions using a centroid-based cosine similarity analysis, comparing validation samples against the archetype centroids established in the discovery cohort (Fig. 3C). This analysis demonstrated that, across all five groups, samples exhibited significantly higher similarity to their assigned archetype than to any alternative subtype, confirming that the classifier successfully identifies samples that are phenotypically congruent with the original discovery definitions. Notably, samples displaying lower cosine similarity relative to their assigned cluster consistently yielded reduced classifier prediction probabilities, suggesting these cases represent biological transition states (Fig. 3C). Finally, a comparison of enrichment profiles between the validation and discovery datasets revealed highly consistent TME phenotypes, further supporting a successful classifier assignment and reproducibility of the derived subtypes (Fig. 3D).

### TME subtypes provide independent prognostic stratification linked to underlying mutational drivers

Next, we evaluated the prognostic stratification of the derived TME subtypes. Kaplan-Meier analysis revealed a significant association between TME subgroups and overall survival (OS), with the “Th/B-cell enriched” group exhibiting the most favorable outcomes and the “Immune-enriched/Fibrotic” group the poorest (Fig. 4A). Multivariate Cox proportional hazards regression confirmed that these TME subtypes are independent prognostic factors (Supp Fig. 4A). Compared to the “Desert” group (reference), the “Th/B-cell enriched” subtype was associated with significantly improved survival (HR = 0.088, p-value = 0.018), while the “Immune-enriched/Fibrotic” group was associated with significantly worse prognosis (HR = 1.970, p-value = 0.042), after adjusting for age, gender, tumor side, Tumor Mutation Burden (TMB), and tumor stage. Notably, within this model, only age and tumor stage remained as significant clinical covariates alongside the TME subtypes.

**Figure 4.**
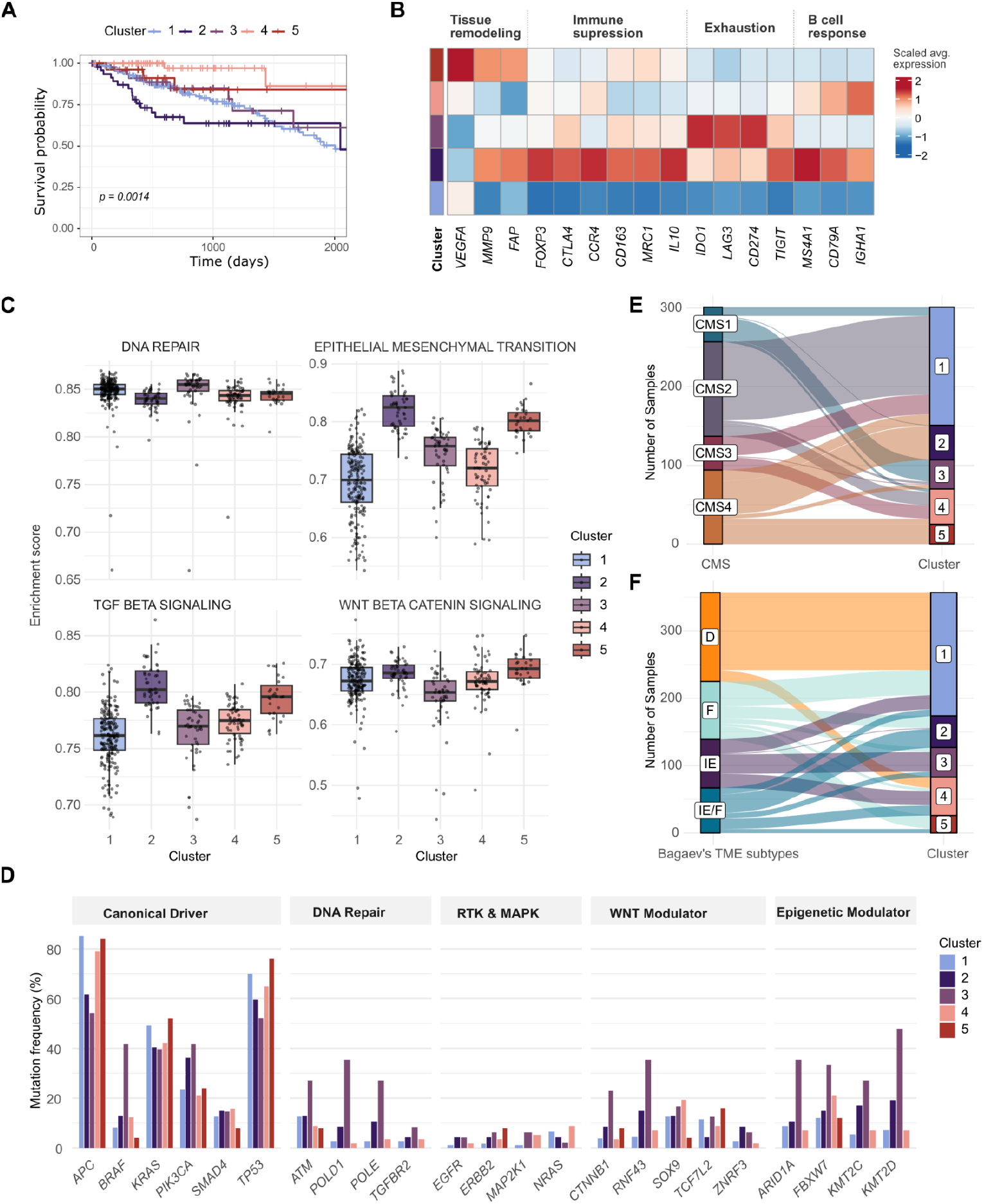
Prognostic stratification and multi-omics characterization of CRC TME archetypes. (**A**) Kaplan-Meier survival curves for overall survival (OS) stratified by the five derived TME subtypes. The “Th/B-cell enriched” group (cluster 4) exhibits superior prognosis, while the “Immune-enriched/Fibrotic” group (cluster 2) shows significantly poorer outcomes. (**B**) Heatmap displaying the average expression of selected immune and stromal modulators. (**C**) Boxplots showing the distribution of enrichment scores for selected MSigDB Hallmark pathways, highlighting subtype-specific biological programs. (**D**) Barplot showing the mutation frequency of known CRC driver genes and key tumorigenic pathways (**E–F**) Concordance with established classifications. Sankey plots mapping the derived TME subtypes to (**E**) Consensus Molecular Subtypes (CMS) and (**F**) Bagaev et al. pan-cancer TME states.

To delineate the biological mechanisms driving these survival differences, we characterized the transcriptional landscape of the subtypes (Fig. 4B + C). Consistent with their enrichment profiles, the two fibrotic groups displayed elevated expression of tissue remodeling markers (Fig. 4B) as well as high TGF-beta signaling and increased expression of genes involved in epithelial mesenchymal transition (EMT) and WNT beta Catenin signaling (Fig. 4C). However, a crucial biological distinction emerged: the “Immune-enriched/Fibrotic” group exhibited high levels of markers for regulatory T cells (*FOXP3, CTLA4, CCR4*) and suppressive M2-like myeloid cells (*CD163, MRC1, IL10*) (Fig. 4B). This profile suggests a highly immunosuppressive microenvironment potentially orchestrated by Cancer-Associated Fibroblasts (CAFs) via TGF-beta signaling, which aligns with the poor prognosis observed in this group (Calon et al. 2015). In contrast, the “Fibrotic/Myeloid” group did not show elevated levels of any immune regulators, likely indicating an immune-excluded phenotype characterized by a physical fibrotic barrier rather than active, cellular-mediated suppression. The “Desert” group displayed a general downregulation of both immune regulation and tissue remodeling programs.

Conversely, the “CD8+ T-cell/Myeloid” group showed the highest expression of checkpoint/exhaustion markers (e.g., *LAG3, IDO1, CD274*) (Fig. 4B) together with high levels of DNA repair genes (Fig. 4C), suggesting an inflamed but exhausted environment that may benefit most from Checkpoint Inhibitor (CI) therapy. The “Th/B-cell enriched” group was defined by genes involved in B-cell responses (Fig. 4B), potentially indicating the presence of Tertiary Lymphoid Structures (TLS), which are known to orchestrate effective anti-tumor immunity and could drive the superior survival observed in these patients (Overacre-Delgoffe et al. 2021; Fridman et al. 2022).

Integration of genomic data revealed distinct mutational profiles underpinning these transcriptomic states. To delineate the genomic landscape of each archetype, we assessed the mutation frequency of known CRC driver genes and key tumorigenic pathways (Fig. 4D). Consistent with an inflamed, hypermutated phenotype, the “CD8+ T-cell/Myeloid” group exhibited the highest tumor mutational burden (TMB) (Supp. Fig. 4B), the highest frequency of Microsatellite Instability (MSI), a large fraction of hypermutated cases (HM-indel and HM-SNV), and an enrichment for right-sided tumors (Supp. Fig. 4C). Accordingly, this subtype displayed the broadest mutational landscape, characterized by significantly elevated mutation frequencies in DNA repair genes (*ATM, POLD1*), epigenetic modulators (*ARID1A, KMT2D*), as well as *BRAF, PIK3CA*, and WNT modulators like *RNF43* and *CTNNB1*. This broad mutational burden and specific involvement of repair mechanisms reinforce its link to the serrated neoplasia pathway and defects in DNA mismatch repair (Leggett and Whitehall 2010; Muzny et al. 2012; Bettington et al. 2013). In stark contrast, the “Desert”,

“Th/B-cell enriched”, and “Fibrotic/Myeloid” groups shared a highly restricted mutational profile dominated by canonical CRC drivers. These subtypes exhibited high mutation frequencies in *APC, TP53*, and to a lesser extent *KRAS*, but maintained relatively low mutation rates across other functional categories. This paucity of broad genomic alterations suggests that these subtypes are primarily driven by canonical WNT signaling within a Chromosomal Instability (CIN) context rather than the hypermutation phenotype observed in the “CD8+ T-cell/Myeloid” group (Muzny et al. 2012; Zhang and Kschischo 2022). The “Immune-enriched/Fibrotic” group displayed an intermediate genomic profile; while it harbored fewer *APC* mutations compared to the CIN-like groups, it showed a modest increase in *PIK3CA* mutations and epigenetic modulators relative to the “Desert” archetype, though these frequencies remained distinctively lower than those observed in the “CD8+ T-cell/Myeloid” group. Collectively, these findings demonstrate that the derived TME archetypes represent distinct clinico-biological entities, linking underlying genomic drivers, such as the serrated neoplasia pathway or chromosomal instability, to specific microenvironmental immune landscapes that ultimately impact patient prognosis.

### TumorArchetypeR refines existing classifications and improves prognostic stratification

Finally, we compared our TME subtypes against existing classification systems. We observed strong concordance with the Consensus Molecular Subtypes (CMS) (Fig. 4E), where tumors are broadly grouped into MSI-Immune (CMS1), Canonical (CMS2), Metabolic (CMS3), and Mesenchymal (CMS4) (Guinney et al. 2015). Specifically, our “Desert” group mapped predominantly to CMS2, the “CD8+ T-cell/Myeloid” group to CMS1, and the “Th/B-cell enriched” group to CMS2/CMS3. Importantly, our model refined the heterogeneity of CMS4—known to be associated with the worst prognosis—by splitting it into two distinct fibrotic subtypes: the “Immune-enriched/Fibrotic” (suppressive) and the “Fibrotic/Myeloid” (potentially excluded). Additionally, the “Th/B-cell enriched” group, which is not explicitly captured by the CMS framework, demonstrated a distinct survival advantage. Together this resulted in superior prognostic stratification by our model compared to CMS (Supp Fig. 4D).

We further compared our results to the pan-cancer TME model proposed by Bagaev et al., which stratifies tumors into Desert (D), Immune-Enriched (IE), Immune-Enriched/Fibrotic (IE/F), and Fibrotic (F) (Bagaev et al. 2021) (Fig. 4F). While general consistency was observed (e.g., Bagaev’s “D” mapping to our “Desert”; “IE” mapping to “CD8+ T-cell/Myeloid”), significant discrepancies highlighted the value of our CRC-specific approach. Bagaev’s “IE/F” and “F” subtypes were not restricted to our fibrotic groups but scattered across multiple subtypes, and the distinct “Th/B-cell enriched” group was entirely missed by the Bagaev classification. Consequently, while the Bagaev “IE/F” group showed a trend toward poorer survival, it failed to reach statistical significance in this cohort (Supp Fig. 4D). In sum, using TumorArchetypeR as a framework to derive TME subtypes from CRC tumors provided a high-resolution dissection of the CRC microenvironment, particularly in resolving fibrotic heterogeneity and identifying a potential TLS-driven phenotype with favorable prognosis.

## Discussion

The successful development of novel cancer therapies, particularly immunomodulatory and stromally targeted agents, relies heavily on dissecting the complex cellular interplay within the TME. However, the integration of TME subtyping into translational research has been hindered by the rigidity of static models, which often fail to capture the specific biological nuances relevant to novel therapeutic mechanisms, and by the technical challenge of applying population-derived signatures to the small, often unbalanced cohorts typical of early-phase trials. In this study, we introduced TumorArchetypeR, a flexible computational framework designed to streamline this translational workflow. By bridging *de novo* subtype discovery with a robust, single-sample classifier, our framework enables the derivation of context-specific archetypes in large discovery cohorts and their consistent propagation to downstream translational datasets. Applying this framework to CRC, we demonstrated its capacity to resolve critical TME heterogeneity that standard classifications miss, thereby providing a rational, data-driven basis for patient stratification to optimize drug discovery processes.

A central biological insight from our analysis is the refinement of the “mesenchymal” poor-prognosis group, broadly categorized as CMS4. Our model resolved this heterogeneous entity into two distinct phenotypes: an “Immune-enriched/Fibrotic” subtype and a “Fibrotic/Myeloid” subtype. While both groups display elevated fibrosis and TGF-beta signaling activity, they differ fundamentally in their immune composition; the “Immune-enriched” variant is defined by the specific recruitment of regulatory T-cells and M2-like macrophages, driving a profoundly immunosuppressive environment associated with the worst overall survival. In contrast, the “Fibrotic/Myeloid” group lacks these active suppressive elements, representing an “immune-excluded” state likely driven by physical barriers. Furthermore, we identified a distinct “Th/B-cell enriched” subtype associated with superior prognosis; the strong B-cell signature in this group is consistent with the potential presence of Tertiary Lymphoid Structures (TLS), which are known to orchestrate effective anti-tumor immunity (Overacre-Delgoffe et al. 2021; Fridman et al. 2022). Crucially, our framework linked these microenvironmental states to distinct cell-intrinsic genomic drivers, suggesting they represent stable biological entities rather than transient states. The “Desert” subtype, alongside the “Th/B-cell” and “Fibrotic/Myeloid” groups, maintained a restricted mutational profile dominated by canonical APC and TP53 mutations, reinforcing the concept of WNT-mediated T-cell exclusion (Luke et al. 2019). In contrast, the inflamed “CD8+ T-cell/Myeloid” subtype mapped strongly to the serrated neoplasia pathway (BRAF, RNF43, POLD1) (Leggett and Whitehall 2010), demonstrating a robust genotype-phenotype concordance that validates the biological stability of these archetypes.

Beyond its application to CRC, the core innovation of TumorArchetypeR lies in its modular and adaptive design, which contrasts with the “black box” nature of many existing subtyping algorithms. By structuring the workflow as a flexible pipeline, we enable users to seamlessly modify, extend, or skip individual processing steps such as scaling, dimensionality reduction, or clustering algorithms thereby tailoring the architecture to specific datasets or biological questions. This flexibility is reinforced by a comprehensive parameter grid search that moves model selection away from arbitrary defaults. By systematically evaluating multiple competing models and providing a suite of stability metrics, covariate associations, and visualization tools, the framework empowers researchers to make informed, data-driven decisions that prioritize biological relevance over mere statistical separation. Furthermore, the integration of a dedicated classifier module specifically addresses the translational gap. By utilizing rank-based binary gene-pair rules that are invariant to cohort-level normalization, the tool ensures that discovered archetypes can be robustly propagated to independent, small, or unbalanced datasets—a critical requirement for clinical implementation. Finally, while we demonstrated its utility in dissecting the TME, the generic nature of this framework suggests it can be readily applied to other complex biological systems, such as classifying tumor-intrinsic molecular states or microbial communities.

### Limitations and Future Directions

Despite these strengths, our study has limitations that highlight important avenues for future research. A notable observation from our classifier results was the identification of samples with intermediate class probabilities. While the classifier’s probabilistic output allows for “soft” assignments that can capture these transitional phenotypes, bulk transcriptomics inherently aggregates signals, making it difficult to distinguish whether these intermediate states represent a true biological transition (e.g., a fibroblast differentiating into a CAF) or simply a spatial admixture of distinct tumor regions. This limitation points to the critical role of spatial context. While not captured in our current data, the flexibility of TumorArchetypeR allows it to ingest diverse feature sets, including cellular neighborhood scores derived from spatial transcriptomics or digital pathology. In particular, leveraging modern machine learning techniques to extract microenvironmental features directly from H&E-stained images offers a promising, clinically accessible route to resolve this spatial heterogeneity. Furthermore, reliance on single-site biopsies captures only a static spatial snapshot of the tumor. In the context of high intra-tumoral heterogeneity, this sampling bias might lead to discordant archetype assignments across different regions of the same lesion, suggesting that future applications should explore multi-region sampling to fully characterize the dominant TME state.

Regarding the specific CRC subtypes identified, it is important to acknowledge that the enrichment scores used for discovery are inferential estimates derived from deconvolution. While scores for major immune lineages showed high concordance across references, the relatively low correlation between malignant cell signatures underscores the challenge of isolating tumor-intrinsic signals from bulk data. Consequently, while we validated the biological logic of these archetypes through genomic and survival associations, orthogonal validation using e.g. single-cell RNA sequencing is necessary to confirm and further resolve the cellular composition and cell-cell interactions (particularly the distinct mechanisms of immune exclusion versus suppression) postulated in our model. Finally, although our classifier demonstrated robustness on a held-out subset of the TCGA cohort, external validation on demographically and technically diverse cohorts will be essential to fully establish the generalizability of the identified CRC archetypes.

## Methods

### PipeComp implementation to derive archetype models

The subtyping workflow is orchestrated via the pipeComp framework (Germain, Sonrel and Robinson 2020), utilizing a PipelineDefinition object to formalize sequential analysis modules (scoring, normalization, scaling, dimensionality reduction, clustering, and characterization). The wrapper function *run_archetype_pipeline* manages dynamic execution, allowing steps to be conditionally skipped based on input metadata. Hyperparameter optimization is achieved by expanding vector arguments (e.g., var_thr, k_neighbors) into a combinatorial grid of competing models. pipeComp handles the parallel execution and caching of these configurations, while dedicated evaluation functions bound to specific steps via stepFn compute stability metrics and clinical associations.

### Implementation of Robust Classifier Training

We implemented the train_classifier module to streamline the generation of rank-based Random Forest models via multiclassPairs (Marzouka and Eriksson 2021). The workflow incorporates a rigorous “consensus filtering” step, which intersects Silhouette scores, k-NN consistency, and Random Forest OOB probabilities to exclude ambiguous or misclassified samples prior to training. The module supports optional class balancing and utilizes k-fold cross-validation to optimize rule-selection hyperparameters. The final production classifier is then trained on the full, curated dataset using these optimized parameters to maximize predictive stability.

### Cell type deconvolution

Cell type abundances were estimated from bulk transcriptome profiles using the xCell2 R package (v1.0.2) (Angel et al. 2025) in R version 4.4.1. We utilized two complementary reference signatures (the Pan-Cancer and TME Compendium collections) obtained from the xCell2 reference repository (https://github.com/dviraran/xCell2refs). The deconvolution algorithm was applied to the discovery and validation datasets independently using default parameters.

### Survival analysis

Survival distributions were estimated using the Kaplan-Meier method and compared via the log-rank test. To assess independent prognostic value, we fitted a multivariate Cox proportional hazards model, adjusting for age, gender, tumor stage, tumor side and TMB. Model validity was confirmed by testing the proportional hazards assumption (Schoenfeld residuals) and checking for multicollinearity (VIF). All analyses were performed using the survival (v.3.8-3) (Therneau 2026) and survminer (v.0.5.1) (Biecek, Kosinski and Biecek 2025) R packages.

### Hallmark enrichment scores

We quantified the activity of known cancer signaling pathways using single-sample Gene Set Enrichment Analysis (ssGSEA) (Barbie et al. 2009). Enrichment scores were derived via the GSVA R package (v.1.52.3) (Hänzelmann, Castelo and Guinney 2013), utilizing the Hallmark gene set collection retrieved from the Molecular Signatures Database (MSigDB v7.4).

### Data Acquisition and Preprocessing

Processed TCGA transcriptomic data was obtained from UCSC Xena Browser (https://xenabrowser.net/). Corresponding somatic mutation profiles were retrieved using the TCGAbiolinks R package (v.2.32.0) (Colaprico et al. 2016), from which Tumor Mutational Burden (TMB) was calculated using the maftools R package (Mayakonda et al. 2018). To benchmark our findings, we utilized pre-existing subtype assignments extracted from their respective original publications, including the Consensus Molecular Subtypes (CMS) (Guinney et al. 2015), Bagaev’s pan-cancer TME states (Bagaev et al. 2021), and genomic classifications (Liu et al. 2018). Tumor purity was quantified using the Consensus Measurement of Purity Estimation (CPE) (Aran, Sirota and Butte 2015), a composite metric derived from the integration of gene expression, DNA methylation, and immunohistochemistry data.

## Code Availability

Code and installation guidelines for the TumorARchetypeR package are available from https://github.com/lutgem/TumorArchetypeR.

---

**Supplementary Figure 1.**
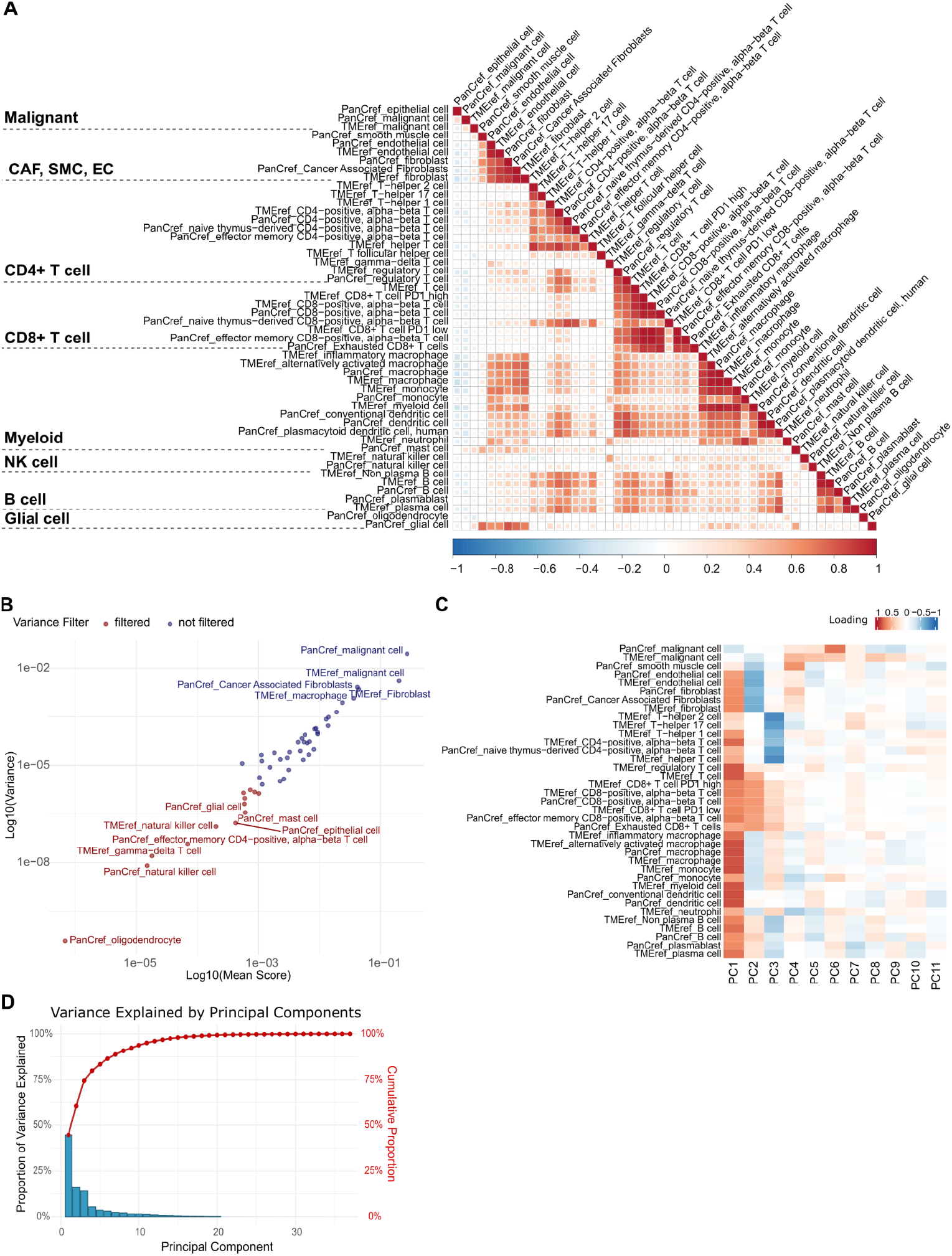
Feature processing and dimensionality reduction of TME signatures. (**A**) Heatmap showing the Pearson correlation of xCell2.0 enrichment scores derived using the Tumor Microenvironment (TMEref) and Pan-Cancer Immune (PanCref) references. The heatmap reveals a high degree of collinearity among features, identifying distinct clusters corresponding to major TME cell lineages. (**B**) Mean-variance relationship of enrichment scores. Scatterplot displaying the mean versus the variance for each cell type signature. Points colored in red indicate low-variance features that were excluded from downstream analysis prior to scaling to prevent the amplification of uninformative noise. (**C**) Heatmap of Principal Component (PC) feature loadings derived from PCA rotation vectors that were scaled by their respective singular values. Loadings represent the correlation between original signatures and the derived components. (**D**) Scree plot illustrating the variance explained by the top Principal Components. Bar heights represent the proportion of variance explained by each individual PC (left y-axis), while the connected red points denote the cumulative variance explained (right y-axis). The cumulative variance is used within the TumorArchetypeR framework to define specific thresholds for selecting the optimal number of PCs to retain for downstream clustering.

**Supplementary Figure 2.**
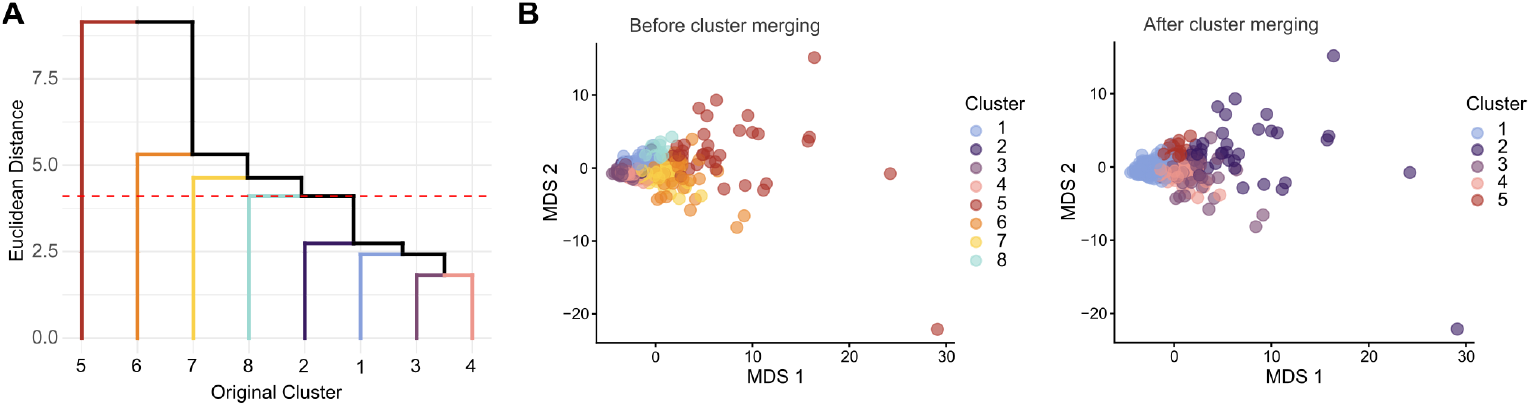
Cluster merging. (**A**) Dendrogram visualizing the hierarchical clustering based on cluster similarity calculated as euclidean distances. The dotted red line marks the cutoff used to merge clusters resulting in a final model with five clusters. (**B**) MDS plots derived from the enrichment score profiles. Each dot represents a sample colored by the cluster assignment before (left panel) or after (right panel) cluster merging.

**Supplementary Figure 3.**
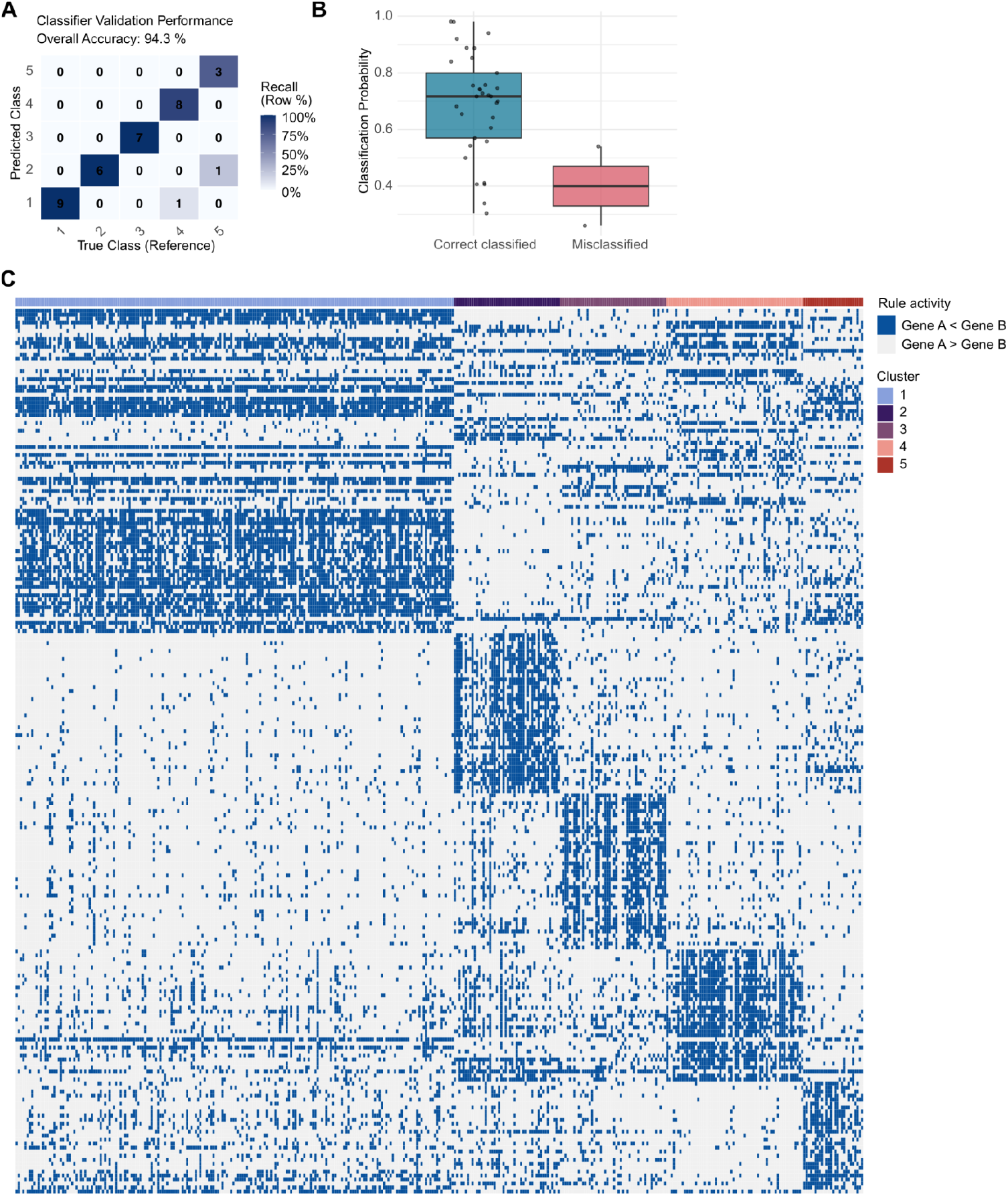
Classifier performance metrics and rule activation patterns. (**A**) Confusion matrix illustrating the training performance of the random forest classifier derived from k-fold cross-validation (k=5) on the discovery dataset (overall accuracy = 94.5%). (**B**) Boxplots comparing the classifier-derived class probabilities for correctly assigned versus falsely assigned samples. (**C**) Binary rule activation landscape. Heatmap displaying the status (active/inactive) of the learned binary gene-pair rules across the training cohort. The clear block-diagonal structure reveals distinct activation patterns for each of the five clusters, confirming that the classifier successfully selected highly discriminative rules specific to each TME subtype.

**Supplementary Figure 4.**
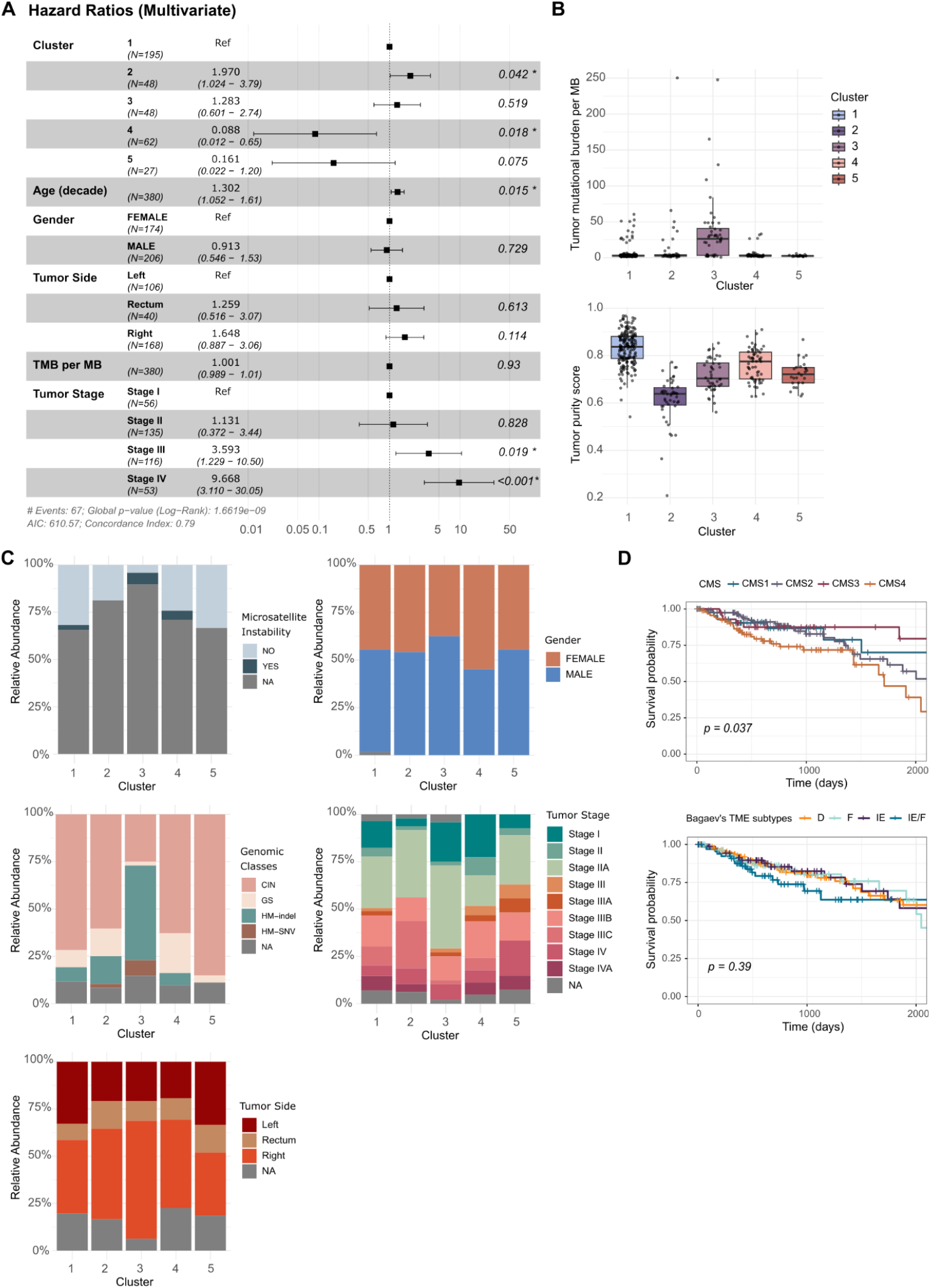
Multivariate survival analysis and associations with covariates. (**A**) Forest plot derived from multivariate Cox proportional hazards regression. The TME subtypes remain independent prognostic factors after adjusting for age, gender, tumor side, TMB, and tumor stage. (**B–C**) Association with genomic and clinical features. (**B**) Boxplots showing the Tumor Mutational Burden (TMB) per MB (upper panel) and tumor purity estimates (lower panel) across subtypes. (**C**) Stacked barplots showing the relative abundance of Microsatellite Instability (MSI) status, gender, genomic class, tumor stage and tumor side across TME subtypes (**D**) Kaplan-Meier plots stratifying the cohort based on CMS (upper panel) and Bagaev et al. (lower panel) classifications.

